# Phage-DMS: a comprehensive method for fine mapping of antibody epitopes

**DOI:** 10.1101/2020.05.11.089342

**Authors:** Meghan E. Garrett, Hannah L. Itell, Katharine H.D. Crawford, Ryan Basom, Jesse D. Bloom, Julie Overbaugh

## Abstract

Understanding the antibody response is critical to developing vaccine and antibody-based therapies and has inspired the recent development of new methods to isolate antibodies. However, methods to define the antibody-antigen interactions that determine specificity or allow escape have not kept pace. We developed Phage-DMS, a method which combines two powerful approaches – immunoprecipitation of phage peptide libraries and deep mutational scanning (DMS) – to enable high-throughput fine mapping of antibody epitopes. As an example, we designed sequences encoding all possible amino acid variants of HIV Envelope to create phage display libraries. Using Phage-DMS, we identified sites of escape predicted using other approaches for four well characterized HIV monoclonal antibodies with known linear epitopes. In some cases, the results of Phage-DMS refined the epitope beyond what was determined in previous studies. This method has the potential to rapidly and comprehensively screen many antibodies in a single experiment to define sites essential for binding to antigen.

## INTRODUCTION

Antibodies are useful research tools, potential therapeutic molecules, and the end goal of many vaccines. Understanding the precise amino acids necessary for binding of antibody to its protein target can provide insights into pathways of escape and improve antigen design for vaccines. Defining these interactions can also enhance our knowledge of antibody function. Modern improvements in the isolation and cloning of monoclonal antibodies (mAbs) have resulted in a dramatic rise in the number of novel antibodies that can be produced, but current methods to map the epitopes of these antibodies cannot presently keep pace. Thus, there is a need for a rapid screening tool to finely map the epitopes of many antibodies in a high-throughput manner.

Structural studies of antibody-antigen complexes are the gold standard for defining key amino acids that directly interact with an antibody, but typically are laborious and require large amounts of antibody. Recently, a method to display libraries of peptides on phage and probe for antibody binding via immunoprecipitation and deep sequencing has been described^1^. This method has been used to map the epitopes of novel HIV-specific mAbs^2-5^, characterize the human virome^6^, and discover autoantigens^7^. Phage libraries offer several advantages over peptide arrays and other mapping methods, namely that phage libraries are easy to generate and store, are relatively low cost, and can be used to rapidly screen for peptide-antibody binding with very small amounts of antibody or plasma. However, while these overlapping peptide libraries are useful for identifying an epitope region, they are limited in their ability to pinpoint individual residues critical for antibody binding.

Several methods exist to more precisely map the specific residues that define an antibody epitope, including the amino acids that disrupt binding and lead to immune escape. Alanine scanning gives single amino acid resolution of antibody epitopes, but it does not provide a complete picture of the potential effect of all possible amino acid mutations at a site. A more comprehensive way to understand the consequences of mutations within the epitope site is to use deep mutational scanning (DMS), which is a technique where each residue of a protein or peptide can be mutated to every possible variant^8^. The resulting library of variants is then used in a functional screen that simultaneously detects the impact of each mutation through deep sequencing. We have previously developed methods employing viral DMS libraries to map the epitopes of HIV-specific antibodies using neutralization as a functional screen^9^. While this approach detects viral escape from neutralization, it is not designed for use with antibodies that bind the viral antigen but mediate their effects through non-neutralizing functions. Therefore, creating a method of mapping antibody epitopes that measures binding agnostic of antibody function is needed.

We have built upon previous studies employing phage display in combination with DMS^10-13^ and here we describe Phage-DMS, a new method that allows high-throughput and high-resolution mapping of antibody epitopes that are proximal in primary sequence with only a single round of immunoprecipitation. Using Phage-DMS, we identified the epitope of four well characterized HIV mAbs, confirming sites of escape predicted using other approaches as well as finding novel epitope sites. Thus, Phage-DMS represents a new tool for the comprehensive mapping of antibody-antigen interactions.

## RESULTS

### Generation and characterization of gp41/V3 HIV Envelope Phage-DMS libraries

A schematic summarizing Phage-DMS is depicted in Figure 1. In brief, we computationally designed sequences that varied at the central position so that this residue contained every possible amino acid corresponding to either the wild type residue or a mutant residue. These sequences were cloned into a phage display vector and then this phage display DMS library was incubated with the antibodies of interest. The resulting phage-antibody complexes were immunoprecipitated and the enriched phage were then subjected to deep sequencing to identify sequences specifically enriched or depleted in the presence of the antibody as compared to the initial phage display library.

**Figure 1.**
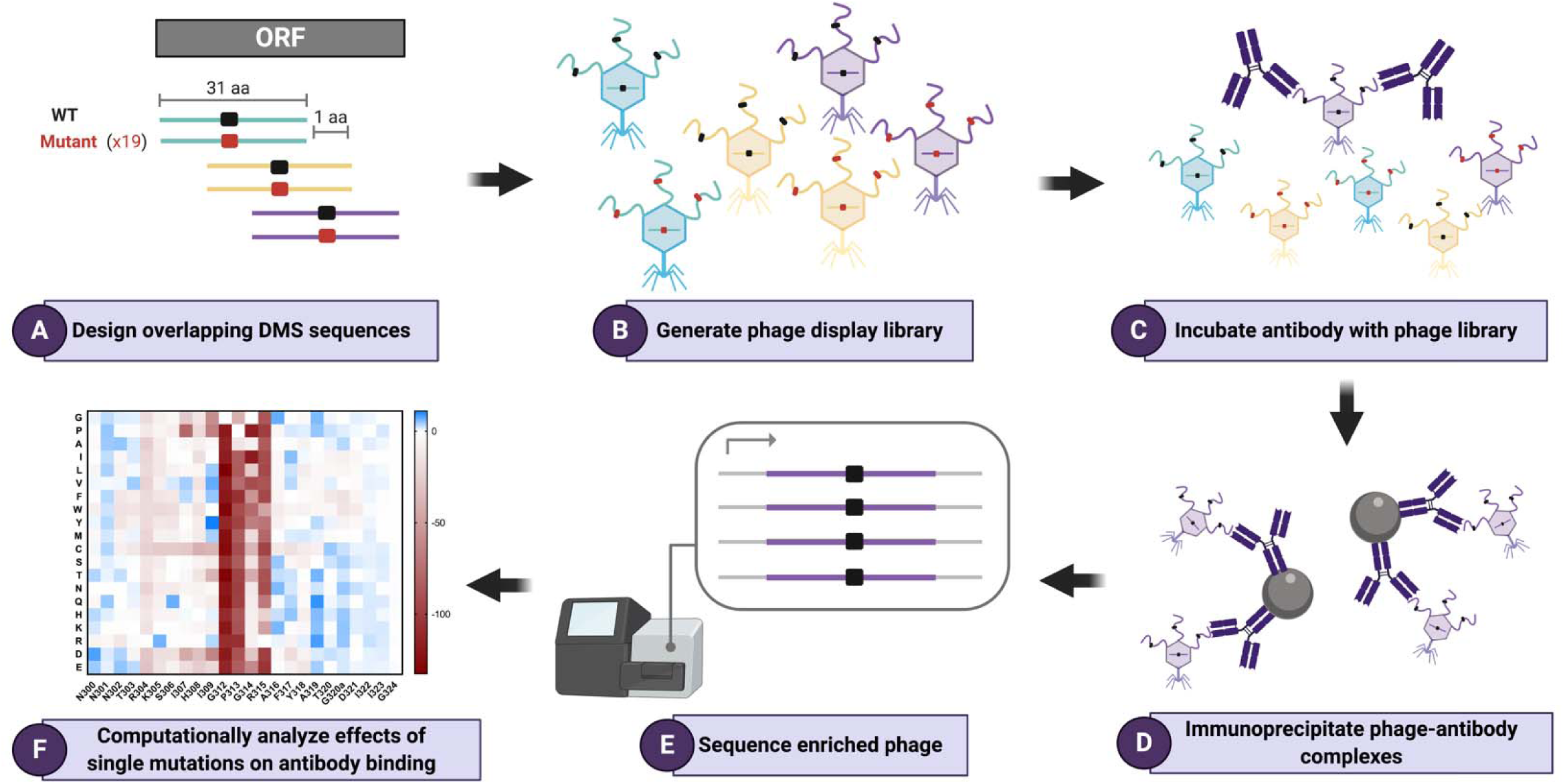
Schematic of Phage-DMS library design and experimental approach. **(A)** To build a Phage-DMS library, sequences are computationally designed to tile across an entire open reading frame of interest, with the central position varying to contain either the wild type residue (shown in black) or a mutant residue (shown in red). **(B)** Sequences are synthesized by releasable DNA microarray and then cloned into a T7 phage display vector. **(C)** The resulting phage display library is incubated with antibody, and **(D)** phage-antibody complexes are immunoprecipitated with magnetic beads. **(E)** Sequences from enriched phage are PCR amplified and pooled samples are deeply sequenced. **(F)** Finally, computational analysis is performed to determine the relative effect of single mutations on the binding of antibody to antigen.

A Phage-DMS library encoding peptides from two immunodominant regions of HIV Env – the V3 region of gp120 and the gp41 ectodomain – was generated in the background of envelope sequences from two common HIV-1 circulating clades^14^: clade A (two variants: BG505, BF520) and clade C (ZA1197) (Supplemental Table 1). A total of 12,160 sequences were included. Duplicate libraries were made starting from the synthesized oligonucleotide array. Fifty-two percent of sequences from gp41/V3 library 1 and 37% of sequences from gp41/V3 library 2 matched the computationally generated sequences, based on sequencing individual plaques (Supplemental Figure 1A). The other plaque sequences contained either indels, point mutations, frameshifts, or multiple errors.

The representation of all expected unique sequences was determined by deep sequencing, performed at a depth of 82- and 45-fold coverage for gp41/V3 library 1 and 2, respectively. A high degree of all unique sequences (91.6% and 87.9%, respectively) were counted at least once in each library (Supplemental Figure 1B and C). There were some regions across gp41 and V3 that were not represented in the libraries, notably sequences corresponding to the fusion peptide domain of gp41. There were also differences in the representation of unique sequences by HIV strain, and we observed that peptides corresponding to BF520 Env were missing more than peptides from other strains. About half (51% and 50%, respectively) of all sequences detected were counted between 10 and 100 times within gp41/V3 library 1 and 2, indicating a modest degree of uniformity (Supplemental Figure 1D).

### Mapping the epitope of gp41- and V3-targeting antibodies with Phage-DMS

To test the ability of our gp41/V3 Phage-DMS replicate libraries to define an antibody epitope, we performed experiments with Env-specific mAbs that have been previously characterized in detail and are known to bind to linear epitopes: gp41-specific mAbs 240D^15^ and F240^16^; V3-specific mAbs 447-52D^17^ and 257D^18^.

#### Gp41-specific mAbs

Deep sequencing of the phage bound by each of the gp41-specific antibodies showed enrichment of wild type sequences corresponding to the expected epitope region of gp41, the immunodominant C-C loop (Figure 2A and B). For both mAbs, peptides in the background of all three Env strains showed enrichment within the same region and to a similar degree. There was a high degree of correlation between technical replicates and good correlation between biological replicates, demonstrating the reproducibility of the Phage-DMS approach (Supplemental Figure 2).

**Figure 2.**
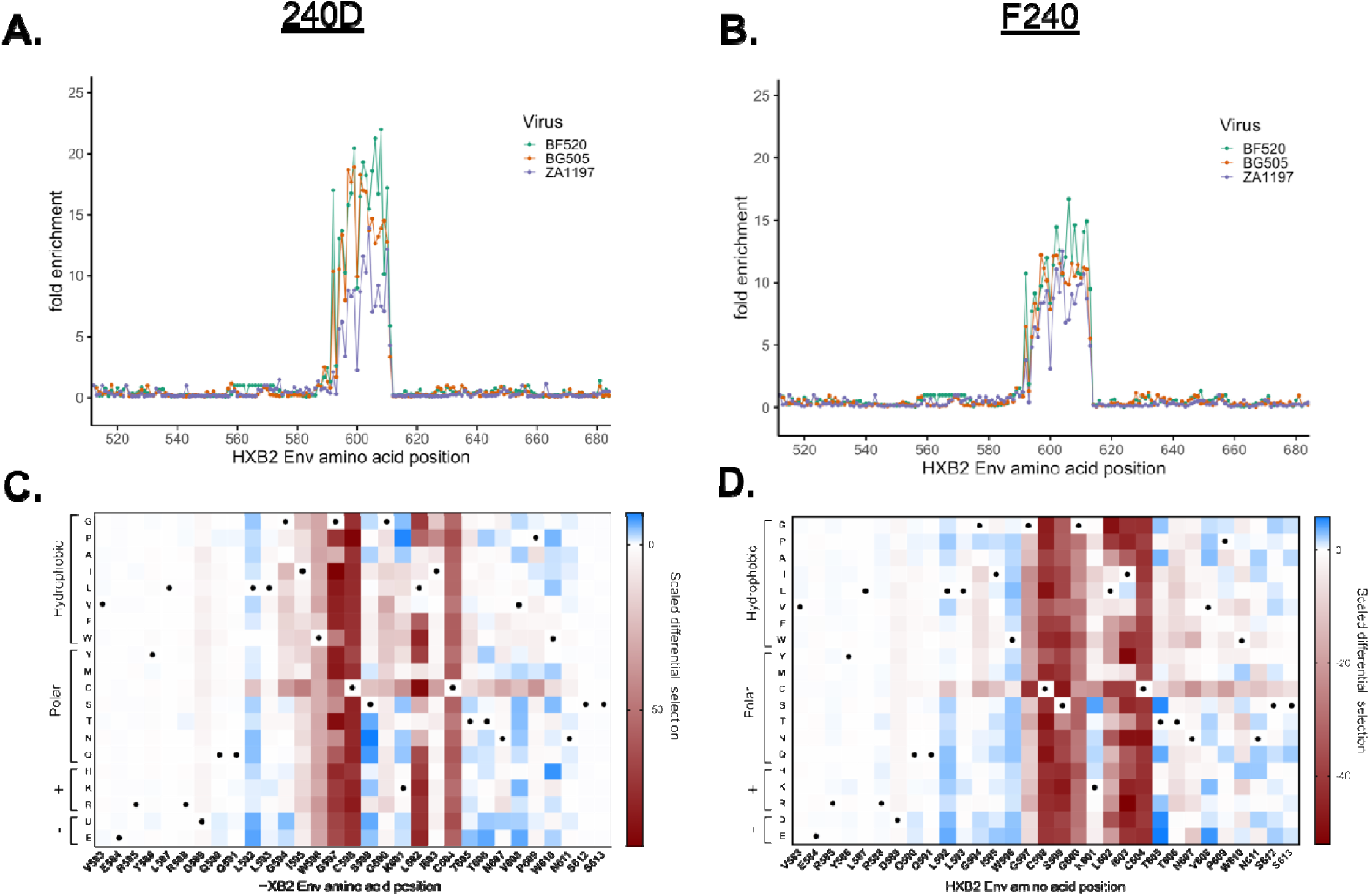
Enrichment and scaled differential selection results from Phage-DMS for gp41-specific mAbs with gp41/V3 libraries. **(A-B)** Line plot showing fold enrichment of wild type peptides in the background of each HIV Env strain for (A) mAb 240D and (B) mAb F240. The color corresponding to each wild type sequence is shown in the upper right. The x axis shows the amino acid position within HIV Env based on HXB2 reference sequence **(C-D)** Heatmap showing the relative effect, as compared to wild type BG505 Env, of each mutation on the binding to (C) mAb 240D and (D) mAb F240, within a selected region of gp41 shown with HXB2 numbering. Wild type residues are marked by a black dot. Amino acids are grouped based on their properties as indicated to the left. All data shown is the average of two biological replicates. See *Quantification and Statistical Analysis* for fold enrichment and scaled differential selection calculations.

To visualize the contribution of each amino acid to the epitope of the antibody, at each site we calculated the differential selection of each mutant-containing peptide as compared to the wild type-containing peptide, scaled by the strength of enrichment of the wild type peptide centered at that site, a metric hereafter termed “scaled differential selection”. Plotting the scaled differential selection for 240D and F240 in the context of the clade A Env protein (BG505) showed strong negative selection at positions C598 and C604 that form the disulfide C-C loop (Figure 2C and D). Mutations to either cysteine residue resulted in very low scaled differential selection values, with no amino acids showing higher preferred binding as compared to wild type, as expected for these C-C loop specific antibodies.

For the 240D antibody, there was also selection against mutations to IWG (aa 595-597; Figure 2C), which was previously identified as the core epitope of 240D^15^. However, mutations to 602 were also negatively selected as compared to wild type, indicating that this site is also important for the epitope. There was enrichment above wild type for most mutations to position L592, as well as single amino acids that are preferred above wild type. For example, the variant N at position 599 and P at position 601 were preferred above wild type, illustrating the potential of this method to identify a sequence that presents a more optimal epitope sequence than the wild type viral sequence.

For the F240 antibody, nearly all mutations in C-C loop residues 597 through 604, with the exception of 601, resulted in depletion in Phage-DMS, suggesting they are part of the epitope (Figure 2D), consistent with structurally-defined interactions for F240^19^. Additionally, some mutations in regions flanking the predicted epitope were positively enriched in Phage-DMS, for example sites 592 and 596.

#### V3-specific mAbs

Phage-DMS with the V3-specific mAbs 447-52D and 257D enriched for wild type peptides corresponding to the expected epitope region of the V3 loop^17, 20^ (Figure 3A and B), with good correlation between replicates (Supplemental Fig 3). For 447-52D, there was binding only to wild type peptides from one of the clade A Env strains (BG505) and the clade C Env strain (ZA1197), but not the other clade A Env strain (BF520) (Figure 3A). This antibody has been reported to bind best to V3 sequences encoding the GPGR motif, but all the Env sequences in this library contain the less preferred GPGQ motif^21^. Sequences outside of this core region differ between the strains that bind (IRIGPGQ) and the strain that does not bind (VHLGPGQ), suggesting these differences may influence binding to 447-52D in the context of the GPGQ motif. There was a similar result with 257D where only BG505 and ZA1197 V3 peptides within the expected epitope region were enriched, again indicating that naturally occurring sequence differences at the tip of the V3 loop may affect binding (Figure 3B).

**Figure 3.**
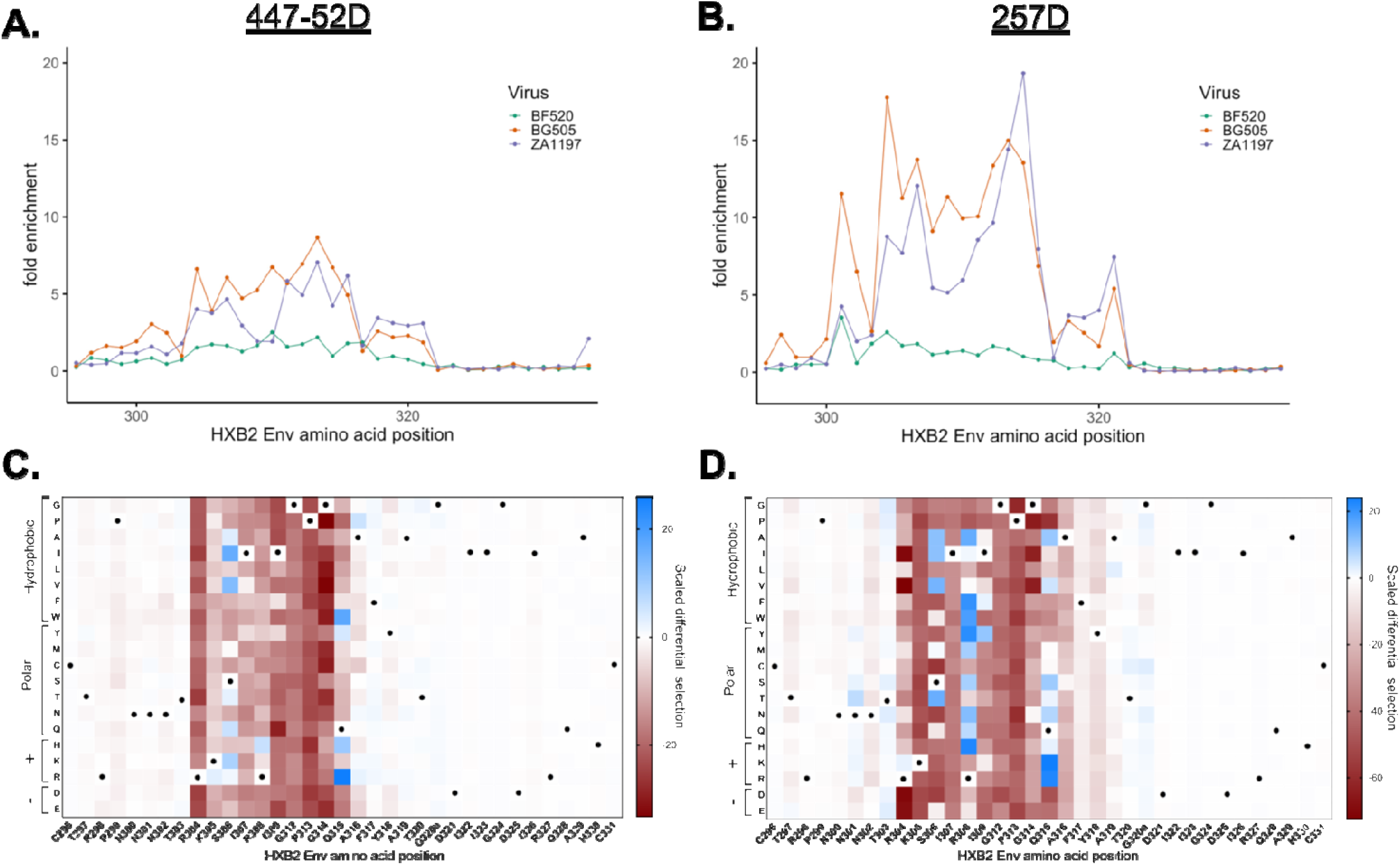
Enrichment and scaled differential selection results from Phage-DMS for V3-specific mAbs with gp41/V3 libraries. **(A-B)** Line plot showing fold enrichment of wild type peptides in the background of each HIV Env strain for (A) mAb 447-52D and (B) mAb 257D. The color corresponding to each wild type sequence is shown in the upper right. The x axis shows the amino acid position within HIV Env based on HXB2 reference sequence **(C-D)** Heatmap showing the relative effect, as compared to wild type BG505 Env, of each mutation on the binding to (C) mAb 447-52D and (D) mAb 257D, within the entire V3 region shown with HXB2 numbering. Wild type residues are marked by a black dot. Amino acids are grouped based on their properties as indicated to the left. All data shown is the average of two biological replicates. See *Quantification and Statistical Analysis* for fold enrichment and scaled differential selection calculations.

The amino acid positions that showed the strongest negative selection with 447-52D in the context of BG505 were in the GPGQ core epitope (Figure 3C). The exception to this is at position 315, where R was the most strongly preferred amino acid. Additionally, variation at sites 304-309 outside of this motif was negatively selected, suggesting these amino acids contribute to the epitope in this context. In particular, mutations to 304 demonstrated consistent negative selection, which has not been reported.

Most mutations to residues spanning 304 to 316 showed strong negative selection with 257D in the context of BG505 (Figure 3D). There were also a number of mutations in this region that demonstrated binding above the wild type residue, including the R at position 315, consistent with the reported preference of 257D for the GPGR motif^22^. H at position 308 was also strongly enriched above the wild type sequence, which agrees with previous studies^20, 22^. Interestingly, there were several other amino acids at position 308 (e. g. F, Y, W) that were more enriched than the wild type R.

### Generation of gp120 HIV Envelope Phage-DMS libraries and validation of V3 epitopes

To test whether we would find similar results with a different Phage-DMS library, a second replicate set of Phage-DMS libraries was constructed, this time containing all peptides spanning the gp120 domain of Env. Given that 9-13% of sequences were missing in the first library, we implemented two changes that were designed to minimize bias against sequences with higher GC content. First, we optimized the GC content of the sequences to be as uniform as possible, with an average GC content of 47%. Second, we implemented subcycling of the annealing and elongation steps between higher and lower temperatures during PCR because this was previously shown to significantly improve amplification of short template pools^23^. 28,840 sequences were generated in the background of a clade A (BG505), clade B (B41), and clade C (DU422) Env and cloned into duplicate phage libraries. Sequencing of individual plaques revealed that 47% of all sequences from gp120 library 1 and 41% of all sequences from gp120 library 2 were correct (Supplemental Figure 4A).

We determined the representation of sequences within each library by deep sequencing at a depth of 71- and 65-fold coverage, respectively. Almost all (96.4% and 96.5%, respectively) unique sequences were present in gp120 library 1 and 2 (Supplemental Figure 4B and C). 66.7% of sequences were counted between 10 and 100 times for gp120 Library 1 and 68.1% for gp120 library 2, demonstrating improved uniformity as compared to the gp41/V3 libraries, where approximately half were represented at this frequency (Supplemental Figure 4D). Our experiments also indicated that subcycling alone improved amplification (Supplemental Figure 4E).

Because the gp120 Phage-DMS libraries we generated contained V3 sequences, we used the same two V3-specific mAbs as above, which enabled the comparison of results between the gp120 libraries and the gp41/V3 libraries. Technical and biological replicates performed with the V3 mAbs and the gp120 Phage-DMS libraries had good correlation (Supplemental Figure 5). When measuring enrichment of wild type sequences for 447-52D, we observed only strong enrichment of wild type peptides from the clade B Env strain, which contains the preferred GPGR motif (aa 312-315) (Figure 4A). 447-52D did not enrich for V3 peptides from the two Env strains that encode GPGQ. For 257D, enrichment of wild type peptides showed binding of this antibody was highest for peptides from the clade B strain, but there was also weak binding to peptides from the clade A strain (Figure 4B). There was no binding of 257D to peptides from the clade C strain.

**Figure 4.**
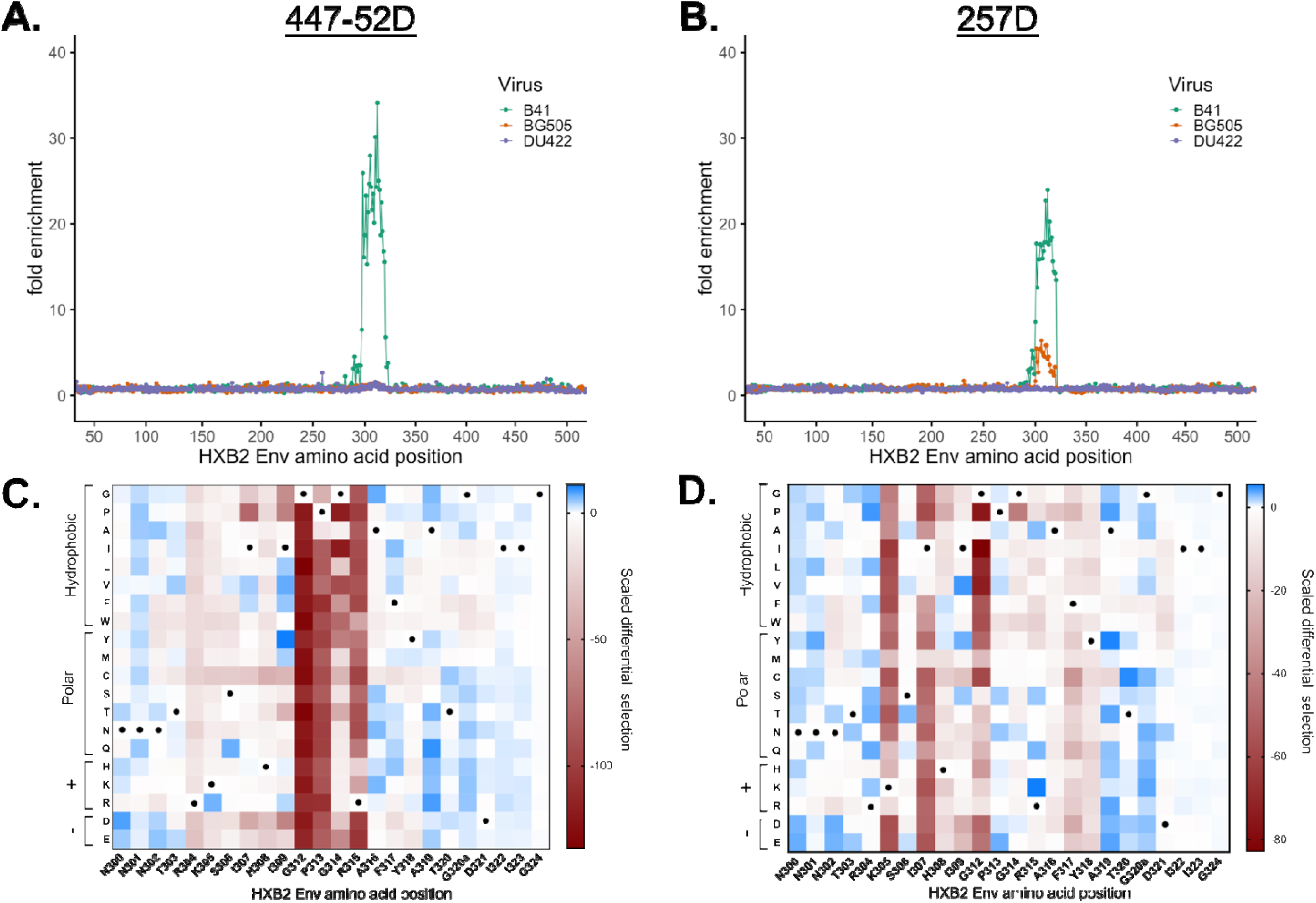
Enrichment and scaled differential selection results from Phage-DMS for V3-specific mAbs with gp120 libraries. **(A-B)** Line plot showing fold enrichment of wild type peptides in the background of each HIV Env strain for (A) mAb 447-52D and (B) mAb 257D. The color corresponding to each wild type sequence is shown in the upper right. The x axis shows the amino acid position within HIV Env based on HXB2 reference sequence **(C-D)** Heatmap showing the relative effect, as compared to wild type B41 Env, of each mutation on the binding to (C) mAb 447-52D and (D) mAb 257D, within a selected region of V3 shown with HXB2 numbering. Wild type residues are marked by a black dot. Amino acids are grouped based on their properties as indicated to the left. All data shown is the average of two biological replicates. See *Quantification and Statistical Analysis* for fold enrichment and scaled differential selection calculations.

Mutations to the residues 312-315 at the GPGR tip resulted in strong negative selection in Phage-DMS experiments with 447-52D (Figure 4C). We additionally saw moderate negative selection at residues upstream of the GPGR, and in particular all mutations to site R304 and H308 resulted in negative selection. For 257D, strongest negative selection was observed for mutations at sites 305, 307, and 312 as compared to wild type (Figure 4D), all of which align with previous studies, which indicated that K-I--GP are the key residues for 257D binding (aa 305-313)^22^. We additionally saw possibly weak contribution of sites on the N-terminal side of the V3 loop (sites 317 and 318) to the epitope.

The observed epitope for both V3 antibodies with the gp120 Phage-DMS libraries was, in general, consistent with previous results with the gp41/V3 libraries. For 447-52D, negative selection at sites 312-315 agreed with previous findings from the gp41/V3 libraries, though the relative strength of negative selection differed between the libraries. For 257D, fewer sites within the epitope region were sensitive to mutation in the context of clade B Env in the gp120 library when compared to results with the gp41/V3 library in the background of clade A Env (BG505). For example, we did not observe that mutations to P313 reduced binding, as we did with the gp41/V3 library, indicating its contribution to the epitope may depend on the context.

### Validation of Phage-DMS results using mutant peptides in a competition ELISA

To confirm that mutations identified by Phage-DMS disrupt binding to the V3- and gp41-specific antibodies, we performed competition enzyme-linked immunosorbent assays (ELISAs) with wild type and mutant V3 or gp41 peptides (see Supplemental Table 2). In this assay, peptides that bind to the antibody block activity against a coated antigen (gp120 or gp41); peptides with mutations in key epitope residues will no longer bind the antibody resulting in an increased IC50 value as compared to a wild type peptide, and peptides that have improved binding to antibody will result in a decreased IC50 value as compared to a wild type peptide.

The wild type gp41 peptide competed for binding of both antibodies to gp41 protein (Supplemental Figure 6A and B). The Phage-DMS results indicated strong preference for the cysteines that form the loop structure for both antibodies, and indeed a gp41 peptide with a mutation in one of these cysteines (C598D) no longer competes for binding to the antibodies at any concentration tested (Figure 5A). Site L602 was also identified as part of the epitope based on a strong depletion of peptides containing mutations to that site. For both 240D and F240, a peptide with a mutation at this site exhibited a 103- and 22-fold increase in IC50, respectively. The IC50 value of peptide S599K was similar to the wild type peptide for 240D but was not detectable for F240, suggesting that the mutant peptide had has lost is ability to bind F240, consistent with the differences observed for these two antibodies by Phage-DMS. A negative control gp41-binding antibody (5F3) did not show reduced activity in the presence of any gp41 peptides (Supplemental Fig 6C).

**Figure 5.**
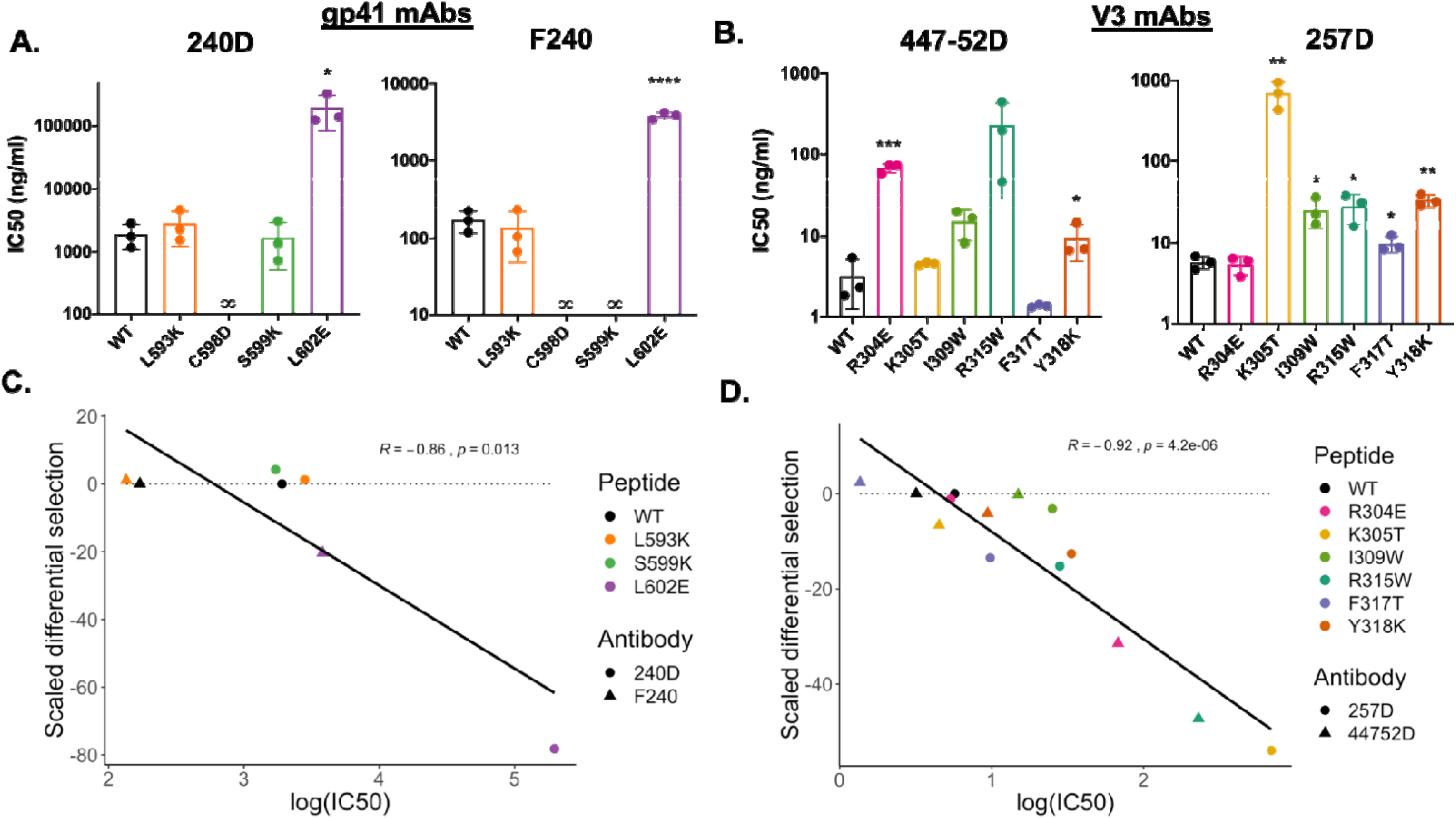
Comparison of effects of mutations on peptide binding in competition ELISAs and Phage-DMS. **(A-B)** Bar plot showing the IC50 values of wild type and mutant peptides in a competition ELISA for (A) gp41-specific mAbs with wells coated with MN gp41 protein or (B) V3-specific mAbs with wells coated with SF162 gp120 protein. Antibodies were pre-incubated with each peptide before addition to the wells. Peptides for which no inhibition of antibody binding could be detected at any concentration tested are indicated with an infinity symbol. Results for three replicate experiments are shown, with the mean +/- SEM. Statistical significance was determined by an unpaired t test. **(C-D)** Correlation between results of IC50 values from competition peptide ELIAs and scaled differential selection values from Phage-DMS for (C) gp41-specific mAbs and (D) V3-specific mAbs. Pearson’s correlation shown at the top.

We confirmed that the wild type V3 peptide competed for binding with both antibodies to gp120 protein (Supplemental Figure 6D and E). With both 447-52D and 257D, the R315W peptide containing a mutation within the important GPGR motif resulted in a 72- and 5-fold increase in IC50, respectively (Figure 5B). Interestingly, the peptide containing a mutation at site 304, which was identified by Phage-DMS as being part of the epitope for mAb 447-52D but not found by any previous studies, demonstrated a 21-fold increase in IC50 as compared to the wild type peptide. Sites 317 and 318 were identified by Phage-DMS as possibly weakly contributing to the epitope, and peptides containing mutations in those sites had a small but statistically significant difference in IC50 from the WT. Peptides with mutations in sites that were not identified by Phage-DMS as being part of the epitope, such as K305T with 447-52D and R304E with 257D, did not yield IC50 values that were significantly different from the wild type peptide in competition ELISAs. VRC01 was used as a negative control gp120-binding antibody and did not show reduced binding in the presence of any V3 peptides (Supplemental Figure 6F).

In order to examine whether the sites predicted to disrupt binding by Phage-DMS correlate with the binding results with corresponding peptides by ELISA, we compared IC50 values obtained for each WT and mutant peptide and differential selection values as measured by Phage-DMS. Differential selection measured for 240D and F240 in the context of the gp41/V3 library and for 447-52D and 257D in the context of the gp120 library each correlated well with the IC50 values measured by competition peptide ELISA (Figure 5C and D), with R^2^ values of 0.74 and 0.85 respectively (p=0.013 and p=4.2e-06).

## DISCUSSION

The process of mapping the epitope of antibodies tends to be labor-intensive with current methods, and often does not provide a detailed understanding of the epitope. To address this, we have combined phage display technology and deep mutational scanning to create an approach called Phage-DMS that is capable of providing a comprehensive view of an antibody epitope in a high-throughput manner. As proof-of-concept, we created Phage-DMS libraries that display wild type and mutant peptides derived from HIV Env strains, and we used these libraries to define the individual amino acids critical for binding to HIV-specific antibodies. The effects of specific mutations identified in this way were validated using the more traditional approach of peptide binding by ELISA. The binding measurements were highly correlated with Phage-DMS scaled differential selection, showing that Phage-DMS measurements capture mutational effects in a relatively quantitative way.

In addition to accurately identifying epitope residues that have been characterized by low-throughput mutagenesis and gold standard structural methods, Phage-DMS also refined and expanded upon the findings from previous studies. For example, mutations to R304 consistently resulted in decreased binding to 447-52D as compared to the wild type sequence, yet this site had not been described to comprise the epitope in previous studies using overlapping hexapeptides^17^ or X-ray crystallography^21, 24^. While neither crystal structure showed electron density at site R304, the prediction from Phage-DMS was validated by competition peptide ELISA, suggesting that Phage-DMS captures information other than physical proximity between antigen and antibody. Conversely, there were instances where previous studies had defined sites on Env with close proximity and hydrogen bonding to the antibody by X-ray crystallography, such as residues 595 to 609 for F240^19^, but we did not observe strong negative selection taking place at sites 595, 596, or 605 - 609. Other studies have demonstrated that side chain proximity does not necessarily predict the contribution of a site to the formation of a protein-protein complex^25-32^, and this case with F240 further illustrates how analysis by Phage-DMS can yield information on the critical binding interactions of an antibody distinct from the identity of interfacing sites obtained by structural studies.

Identification of an antibody’s epitope by Phage-DMS was reproducible, as demonstrated by the consistent results obtained by testing the same antibodies with two independently generated libraries. However, we did observe differences in the results with libraries built with different viral strains, highlighting the sensitivity of this assay to viral context. This occurred in the case of 447-52D, where one library had a viral strain with a more optimal core epitope (B41 in the gp120 library), but the other library (gp41/V3) did not. In this case, other sites sensitive to mutation were revealed in the library where none of the wild type sequences contained the optimal core motif. Phage-DMS has several advantages over other mapping approaches. The high titer and renewable nature of phage libraries make them ideal for large-scale and high-throughput experiments. Antibodies can be tested against multiple background sequences in parallel, which is especially important when studying antibodies against a pathogen as diverse as HIV. Other mutation-based approaches often involve creating and testing mutants in the background of a single Env strain. Phage-DMS can identify the binding sites for neutralizing antibodies, like 447-52D and 257D, and non-neutralizing antibodies, like 240D and F240, whereas many other methods usually rely on a functional read out, such as the ability to escape neutralization. Moreover, unlike methods that involve growth of the virus in culture, phage will display peptides encoding mutations irrespective of their effect on replication fitness, resulting in a more complete picture of the potential effect of all variants on binding. Given its versatility, Phage-DMS is not limited to mapping antibody epitopes and could theoretically be used to map binding sites between any two proteins of interest, similar to other studies that have utilized DMS to dissect protein-protein interactions using phage^10-13^.

There are some limitations of Phage-DMS. For one, post-translational modifications such as glycosylation are not present on the peptides displayed by the phage. Additionally, due to the length of peptides displayed in these libraries, Phage-DMS in its current form is limited to mapping epitopes that dependent on proximal amino acids and thus is not suited for complex conformational epitopes. To confirm these limitations, we tested known glycan-dependent (BF520.1, QA013.2) and discontinuous epitope (PGT145, 50-69) HIV Env-specific mAbs in Phage-DMS experiments, and as expected did not observe enrichment of any peptide sequences (data not shown). However, our data and findings by others who have generated phage display libraries suggest that peptides can fold and present native-like conformations to some extent^7^. For example, Phage-DMS showed a strong preference for cysteines that form a predicted loop structure within the epitope region of the gp41-specific antibodies.

Phage-DMS can be applied not just towards identifying epitope sites and potential pathways of antibody escape, but also to improving the binding of antibody to antigen. We observed cases where mutations, such as F317T with antibody 447-52D, improved binding above wild type levels by both Phage-DMS and competition peptide ELISA, demonstrating that Phage-DMS results could optimize antigen targeting and aid in rational vaccine design. Overall, we have demonstrated that Phage-DMS can map sites essential for antibody binding and could potentially accelerate the characterization of mAbs and development of vaccines.

## METHODS

### Antibody production

The following antibodies were obtained from the AIDS Reagent Program, Division of AIDS, NIAID: 5F3, F240, 240D, VRC01, 257D, 447-52D.

### Generation of Phage-DMS libraries

HIV-1 strains included in the gp41/V3 DMS phage library and gp120 DMS phage library are noted in Supplemental Table 1. We did not include the signal peptide of gp120 and the transmembrane and cytoplasmic domains of gp41 in these libraries. Oligonucleotides coding for peptides 31 amino acids in length were computationally designed to overlap by 30 amino acids and tile across the entire length of these protein sequences. Sequence coding for a linker ([G_4_S]_3_ for gp120 and gp41 sequences, and 15 flanking wild type residues for the V3 sequences) was added to the beginning and end of the protein sequence of interest in order to ensure that every residue of the protein was located at the central position of a peptide ^33^. All sequences were codon optimized for expression in *E. coli* by either using IDT online Codon Optimization Tool (http://www.idtdna.com/CodonOpt) for the gp41/V3 libraries or the Codon Optimization OnLine tool (COOL, http://cool.syncti.org/) for the gp120 libraries. Linker and protein coding sequences for the gp120 DMS phage library were additionally GC optimized using the COOL tool, with a target GC concentration of 50%. The central codon of each oligo was replaced with codons representing every possible amino acid, generating a total of 20 oligos centered at each site along the protein. Each oligo additionally had 5’ and 3’ adaptor sequences added to facilitate amplification and cloning (5’: AGGAATTCTACGCTGAGT, 3’: TGATAGCAAGCTTGCC). After de-duplicating oligonucleotides that were identical across the different HIV strains, 12,160 unique sequences for the gp41/V3 library and 28,840 unique sequences for the gp120 library were synthesized as an oligonucleotide pool by Twist Bioscience. Code used to generate oligonucleotide sequences can be found at github.com/XXX.

Oligonucleotide pools were resuspended in Tris-EDTA (TE) buffer, pH 8.0, to a concentration of 10 ng/uL. PCR amplification of the pools with the KAPA HiFi PCR Kit (Roche) was done with 2.5 uL of a 2 ng/uL solution of the pool in a 50uL reaction, with primers annealing to the adaptor sequences (Fwd: AATGATACGGCAGGAATTCCGCTGAGT, Rvs: CGATCAGCAGAGGCAAGCTTGCTATCA). Amplification of the gp41/V3 oligos was performed using the following thermocycler conditions:

1. 95°C for 3 min
2. 98°C for 20 s
3. 60°C for 15 s
4. 72°C for 15 s
5. **Go to step 2 (x3)**
6. 95°C for 30 s
7. 98°C for 20 s
8. 72°C for 30 s
9. **Go to step 7 (x20)**
10. 72°C for 5 min
11. Hold at 4°C

Amplification of the gp120 library was performed including subcycling steps (adapted from ^23^), using the following thermocycler conditions:

1. 95°C for 3 min
2. 98°C for 20 s
3. 60°C for 15 s
4. 72°C for 15 s
5. **Go to step 2 (x3)**
6. 95°C for 30 s
7. 98°C for 20 s
8. 67°C for 15 s
9. 72°C for 15 s
10. **Go to step 8 (x4)**
11. **Go to step 7 (x20)**
12. 72°C for 5 min
13. Hold at 4°C

PCR products were cleaned using Agencourt AMPure XP beads (Beckman Coulter) and then digested with EcoRI-HF and HindIII-HF (NEB) at 37°C for 1 hour. Cloning into bacteriophage was performed using the T7Select System (EMD Millipore). In brief, four separate digests of the PCR products were performed and then pooled, purified using gel electrophoresis, and ligated at a 3:1 vector to insert ratio into the T7Select 10-3b bacteriophage vector arms overnight at 16°C. 5 uL of the ligation reaction was added 25 uL of T7Select Packaging Extract and incubated at room temperature for 2 hours, then stopped using 270 uL sterile LB. The phage titer from the packaging reaction was determined using a plaque assay, then amplified on the host E coli (BLT5403) and titered again. At every step each library member was represented by >200 plaque forming units (pfu) to avoid bottlenecking, and the final libraries had >200,000 pfu/mL per unique library member. Sanger sequencing of at least 40 plaques from each library was performed to assess the diversity and fidelity of the phage libraries. This process was performed in duplicate, starting from the PCR amplification step, and the final libraries were stored at -80°C with 0.1 volumes sterile 80% glycerol and penicillin/streptomycin added.

### Antibody immunoprecipitation with Phage-DMS library

Immunoprecipitation with antibody bound to peptide displayed on phage was performed in a manner previously described^1, 2^. In brief, the phage library was first thawed and diluted to a concentration representing 200,000 pfu/mL per unique library member (for example, the gp120 DMS phage library has 28,840 members, therefore we use a concentration of 5.8 × 10^9^ pfu/mL). After blocking a deep 96 well plate with 3% BSA in TBST, 1 mL of the diluted phage library with 10 ng of the antibody of interest was added to the well. Plates were sealed and incubated on a rocker at 4°C for 20 hours. 20 uL of each Protein A and Protein G Dynabeads (ThermoFisher) were added to each well and incubated on a rocker at 4°C for 4 hours. Magnetic separation was performed and antibody-bead complexes were washed 3x with 400 uL wash buffer (150 mM NaCl, 50 mM Tris-HCl, 0.1% [vol/vol] NP-40, pH 7.5). Beads were resuspended in 40 uL of water, transferred to a 96 well PCR plate and then bound phage lysed at 65°C for 10 minutes. Antibody selections were done in technical duplicate, and duplicate mock selected samples were included in order to determine the background levels of peptide binding to the beads. Additionally, we lysed 10-20 million phage from the diluted input library to determine the distribution of phage in the starting library.

### Deep sequencing

To determine the frequency of each peptide in the antibody selected, mock selected, and input conditions, we deep sequenced the lysed phage from each sample. We performed two rounds of PCR to amplify and add dual barcodes to each sequence. Each PCR reaction was performed using Q5 Hot Start High Fidelity 2X Master Mix (NEB). For the first round of PCR, 20uL of lysed phage was used as the template in a 50 uL reaction along with primers that anneal to sequences on either side of the cloning region within the T7 Select 10-3b vector (Fwd: TCGTCGGCAGCGTCTCCAGTCAGGTGTGATGCTC, Rvs: GTGGGCTCGGAGATGTGTATAAGAGACAGCAAGACCCGTTTAGAGGCCC). We used the following thermocycler conditions for the first round of PCR:

1. 98°C for 30 s
2. 98°C for 5 s
3. 68°C for 10 s
4. 72°C for 30 s
5. **Go to step 2 (x30)**
6. 72°C for 2 min
7. Hold at 4°C

2 uL of the PCR product was then used as the template in a 50 uL reaction for the second round of PCR, with indexed P5 and P7 primers that anneal to sequence added by the round 1 primers (Fwd: AATGATACGGCGACCACCGAGATCTACACNNNNNNNNTCGTCGGCAGCGTCTCCAGTC, Rvs: CAAGCAGAAGACGGCATACGAGATNNNNNNNNGTCTCGTGGGCTCGGAGATGTGTATAAGAGA CAG, where “NNNNNNNN” denotes a unique 7-nt indexing sequence). We used the following thermocycler conditions for the second round of PCR:

1. 98°C for 30 s
2. 98°C for 5 s
3. 72°C for 40 s
4. **Go to step 2 (x8)**
5. 72°C for 2 min
6. Hold at 4°C

PCR products were then cleaned using Agencourt AMPure XP beads and eluted in 50 uL water. DNA concentrations were quantified via Quant-iT PicoGreen dsDNA Assay Kit (ThermoFisher). Equal amounts of DNA from the antibody selected samples were pooled, along with equal amounts of mock selected and input library at 10X the amount of the antibody selected samples (for example, 20 ng of each antibody selected sample was pooled along with 200 ng of each mock selected sample and input library sample). Finally, the pooled sample was gel purified, quantified using the KAPA Library Quantification Kit (Roche), and then sequenced on an Illumina MiSeq with 1×125 bp single end reads using a primer that anneals to the 5’ adaptor sequence just prior to the oligo sequence (GCTCGGGGATCCGAATTCTACGCTGAGT).

### Competition peptide ELISA

Custom peptides were synthesized by ThermoFisher. V3 peptides were resuspended in water at 1 mg/mL, and gp41 peptides were resuspended in 50% DMF due to increased hydrophobicity and to prevent degradation of the cysteine disulfide bond. To perform a competition peptide ELISA, plates were first coated overnight with 500 ng/mL of either SF162 gp120 (Cambridge Biologics, # 01-01-1063) or MN gp41 (AIDS Reagent Program, # 12027) in PBS. After washing 4X with wash buffer (1X PBS, 0.05% Tween-20), wells were blocked with blocking buffer (1% BSA in PBS + 1mM EDTA) for 1 hour at 37°C. Antibodies diluted to 500 ng/mL in blocking buffer were preincubated for 30 minutes with peptide at various concentrations, with controls incubated with equivalent volume of appropriate peptide resuspension buffer to establish baseline activity. Blocking solution was washed from plates and then antibody-peptide samples were added to each well and the plate was incubated for 1 hour at 37°C. Plates were washed and then goat anti-human HRP (Sigma) diluted 1:2500 in blocking buffer was added to each well and incubated for 1 hour at 37°C. After washing again, 1-Step Ultra TMB-ELISA Substrate Solution (ThermoFisher) was added and incubated at room temperature for either 15 minutes (V3 peptides) or 5 minutes (gp41 peptides) before adding an equivalent volume of 1N H_2_SO_4_. OD_450_ value for each well was then measured on a plate reader. All assays were performed in 384 well plates (ThermoFisher) and washed by robot. IC50 values are calculated by taking the OD_450_ values across a dose response curve of the peptide and computing in Prism (GraphPad version 8.4.0) the concentration of peptide which inhibits half of the maximum activity (e.g. activity with no peptide present).

## Quantification and Statistical Analyses

Code used to perform the following analyses and generate plots with Phage-DMS data can be found at https://github.com/meghangarrett/Phage-DMS

### Demultiplexing and alignment

Demulitplexing and fastq file generation were performed using Illumina MiSeq Reporter software. Reads were aligned to their respective reference libraries using bowtie v1.1.1^34^, with the options “--trim3 32 -n 0 -l 93 --tryhard --nomaqround --norc --best --sam –quiet”. We aimed to get 10X coverage of antibody selected samples and 100X coverage of mock selected samples and input library, keeping in mind that only about half of the phage in the library contained sequences that perfectly matched the computationally designed sequences. All reads were deposited to NCBI and are accessible under BioProject # PRJNA630833.

### Calculating enrichment of peptides

The enrichment (*E*_*r,x*_) of a peptide containing amino acid *x* centered at site *r* is calculated as follows. First, read counts *n* from technical replicates are added together. The proportion *p* of each peptide within a sample (Equation 1) or within the input library (Equation 2) is calculated by comparing the read count of each peptide with amino acid *x* at site *r* to the sum of all read counts in the sample.

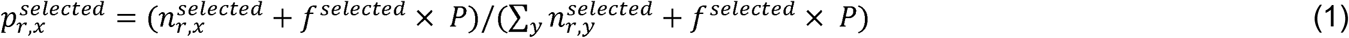

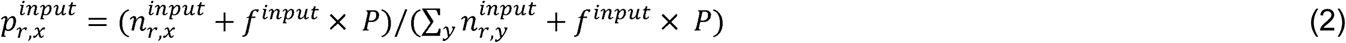

To account for statistical noise, a pseudocount of *P*=1 is added to each count. To scale the pseudocount according to the varying sequencing depth of the antibody selected sample and library input, we calculated *f*^*selected*^ (Equation 3) and *f*^*input*^ (Equation 4) variables as follows.

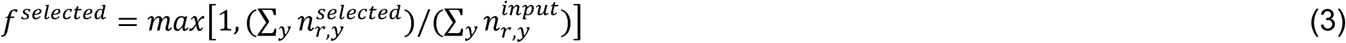

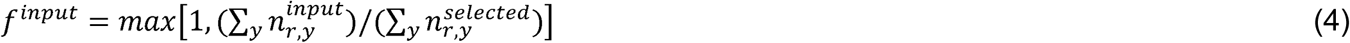

To calculate the enrichment *E*_*r,x*_, the proportion 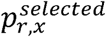 of each peptide within the antibody selected sample is compared against the proportion 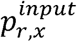of each peptide within the input library (Equation 5)

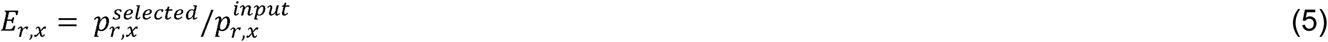

### Calculating differential selection and scaled differential selection

Differential selection is a metric used to calculate the relative enrichment of a peptide containing a mutation (*E*_*r,x*_) as compared to the enrichment of a peptide containing the wild type amino acid at a site (*E*_*r,wt(r)*_) (Equation 6).

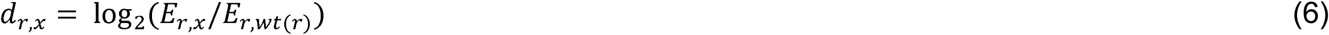

By definition, the differential selection of the wild type amino acid at a site (*d*_*r,wt(r)*_) is always 0.

In order to emphasize the differential selection taking place at sites within the epitope, we calculated a metric we termed “scaled differential selection”. To get the scaled differential selection *S*_*r,x*_ of a mutation *x* at a site of interest *r*, we take the differential selection as calculated above and multiply it by the enrichment of the peptide containing the wild type amino acid centered at the position of interest (Equation 7).

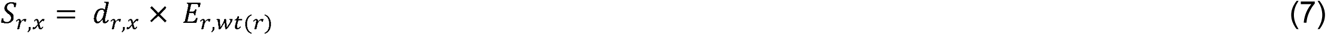

## ACKNOWLEDGEMENTS

We thank T. Gobillot and A. Dingens for helpful discussions, C. Sather and the Fred Hutch Genomics Core for assistance with sequencing, and L. Stamatatos for generous use of lab equipment. This work was supported by NIH R01 AI138709 and AI120961. MEG was supported in part by T32 AI083203.

## CONTRIBUTIONS

JO conceived the project; MEG and JO led the design of the study; MEG and HLI contributed to experimental design and performed experiments; MEG, KDC, RB and JDB participated in computational and data analyses. MEG and JO wrote the paper with input from all authors.

## FIGURES

**Supplemental Table 1.** Description of strains use to construct Phage-DMS libraries. Listed are the strains of HIV Env that were used to generate tiling wild type or mutant sequences for either the gp41/V3 or gp120 Phage-DMS libraries.

| Common name | Strain | Codon optimized? | GC optimized? | Source | GenBank # | Env AA sites included in library (autologous numbering) |
| --- | --- | --- | --- | --- | --- | --- |
| **Gp41/V3 Phage-DMS library** |  |  |  |  |  |  |
| BG505 | BG505.W6M.ENV.C2 | yes | no | Xu et al | DQ208458.1 | 295 - 329, 509 - 681 |
| BF520 | BF520.W14M.C2 | yes | no | Simonich et al | KX168094.1 | 290 - 324, 501 - 673 |
| ZA1197 | C.ZA.1197MB | yes | no | Rousseau et al | AY463234.1 | 293 - 327, 494 - 666 |
| **Gp120 Phage-DMS library** |  |  |  |  |  |  |
| BG505 | BG505.W6M.ENV.C2 | yes | yes | Xu et al | DQ208458.1 | 30 - 508 |
| B41 | 9032-08.A1.4685 | yes | yes | Keele et al | EU576114.1 | 30 - 515 |
| DU422 | Du422.1 | yes | yes | Li et al | DQ411854.1 | 30 - 506 |

**Supplemental Table 2.**
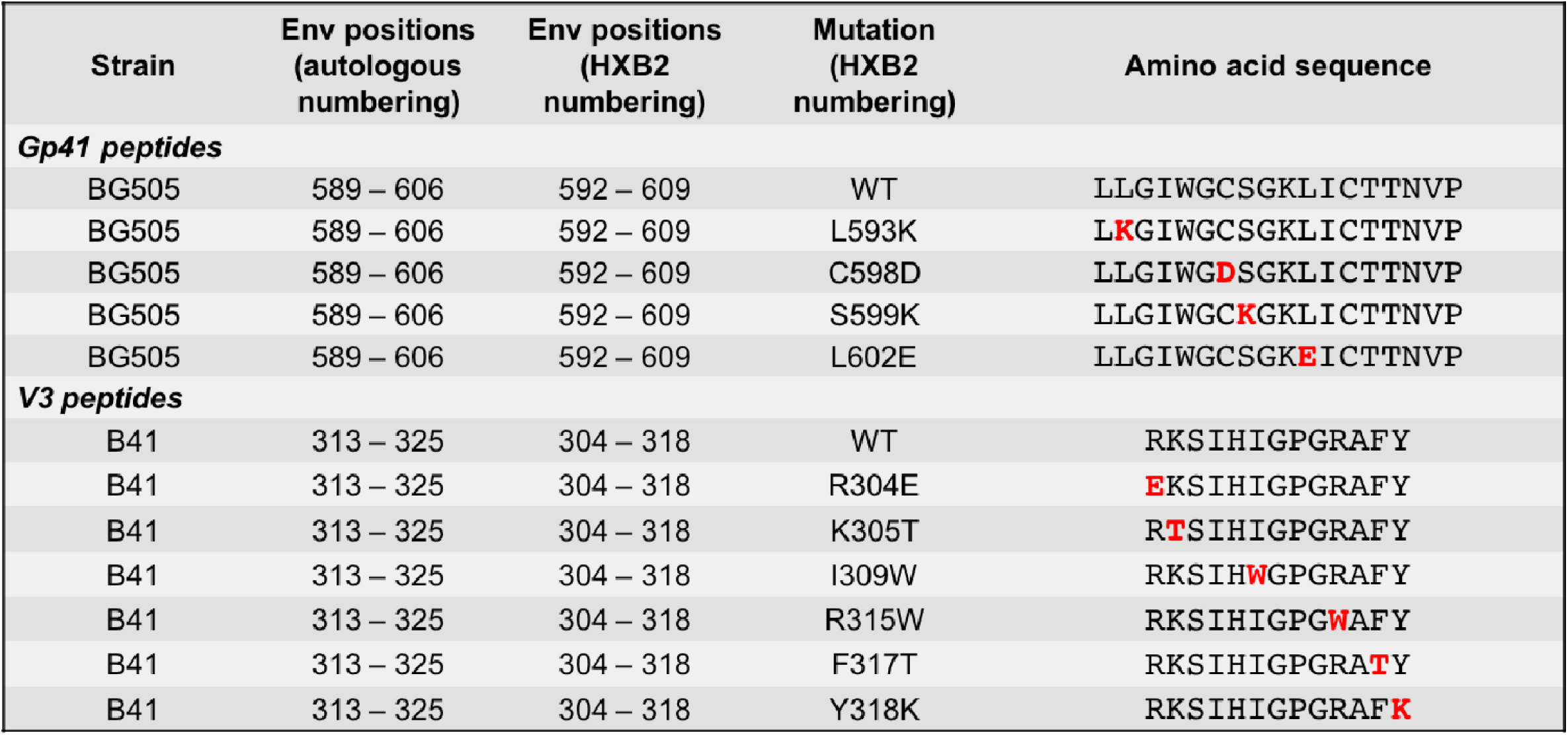
Description of peptides used to perform competition ELISAs. Mutations are highlighted in red.

**Supplemental Figure 1.**
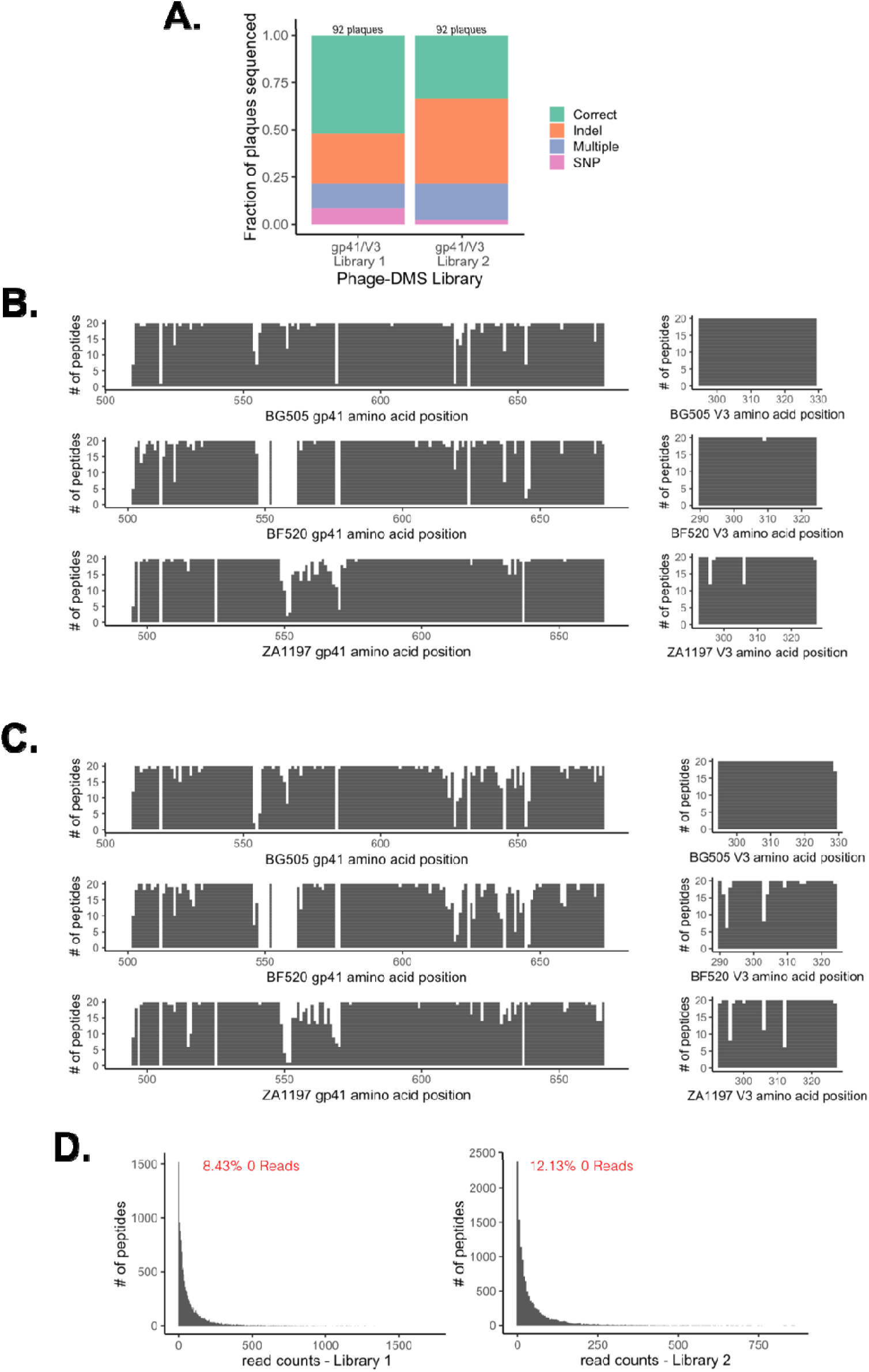
Composition and distribution of sequences in the gp41/V3 Phage-DMS libraries. **(A)** Stacked bar plot depicting results of Sanger sequencing individual plaques from each library. Total number of plaques sequenced is displayed at the top. SNP = single nucleotide polymorphism. **(B)** Coverage of all unique sequences centered at sites across gp41 and V3 within gp41/V3 library 1, as determined by deep sequencing at 82-fold coverage. The height of the line at each site corresponds to whether each unique amino acid variant was counted at least once, with 20 total possible amino acids. **(C)** Coverage of all unique sequences centered at sites across gp41 and V3 within gp41/V3 library 2, as determined by deep sequencing at 45-fold coverage. **(D)** Histogram showing the distribution of all reads sequenced from gp41/V3 library 1 and 2 as determined by deep sequencing. Percent of computationally designed sequences not counted shown in red.

**Supplemental Figure 2.**
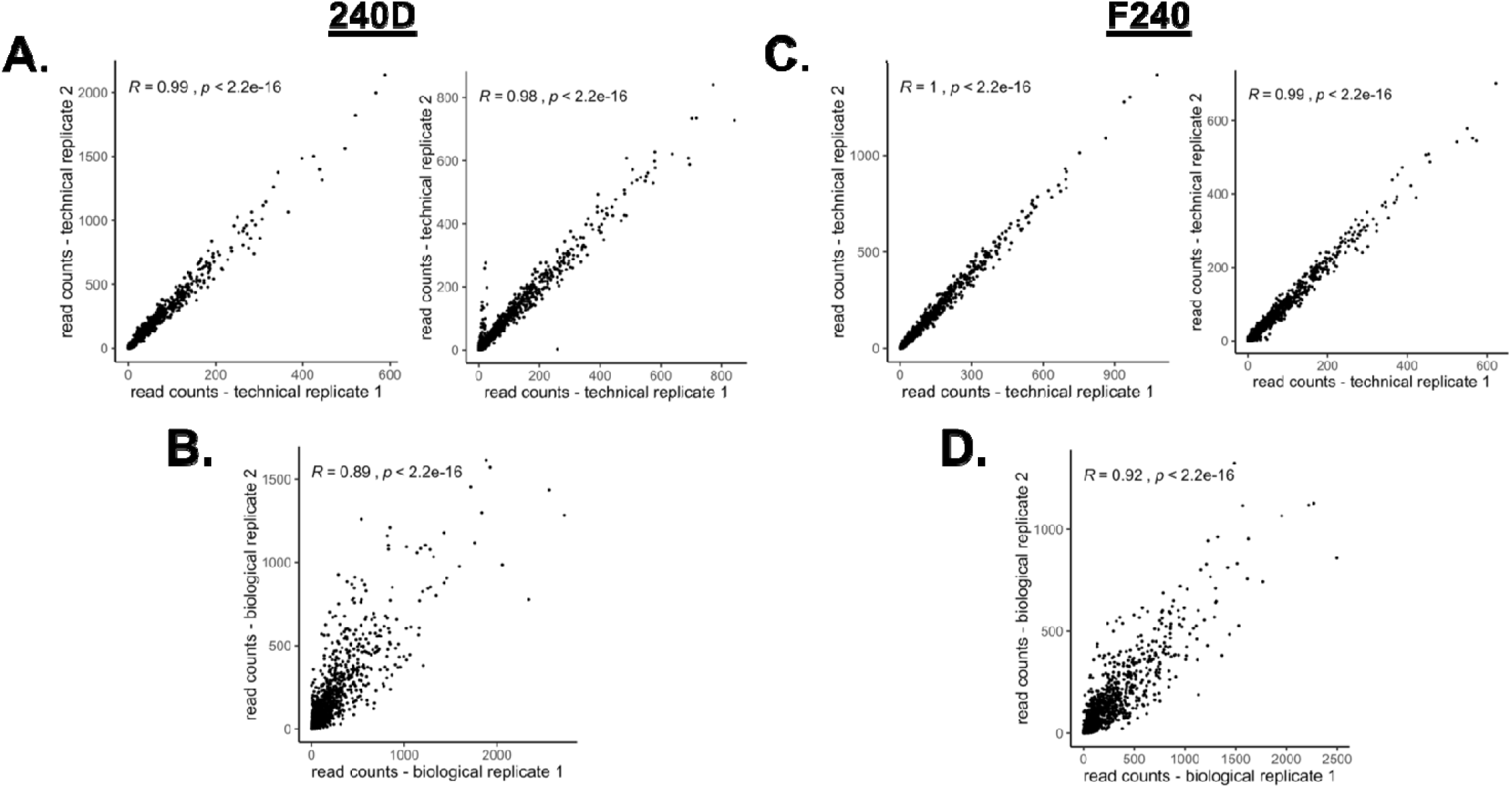
Reproducibility of gp41/V3 Phage-DMS experiments with gp41-specific antibodies. Correlation of raw read counts between technical and biological replicates for mAb 240D and mAb F240. **(A, C)** Correlation between replicate wells done in parallel for each experiment. **(B, D)** Correlation between separate experiments performed with gp41/V3 library 1 and 2. Pearson’s correlation is shown at the top.

**Supplemental Figure 3.**
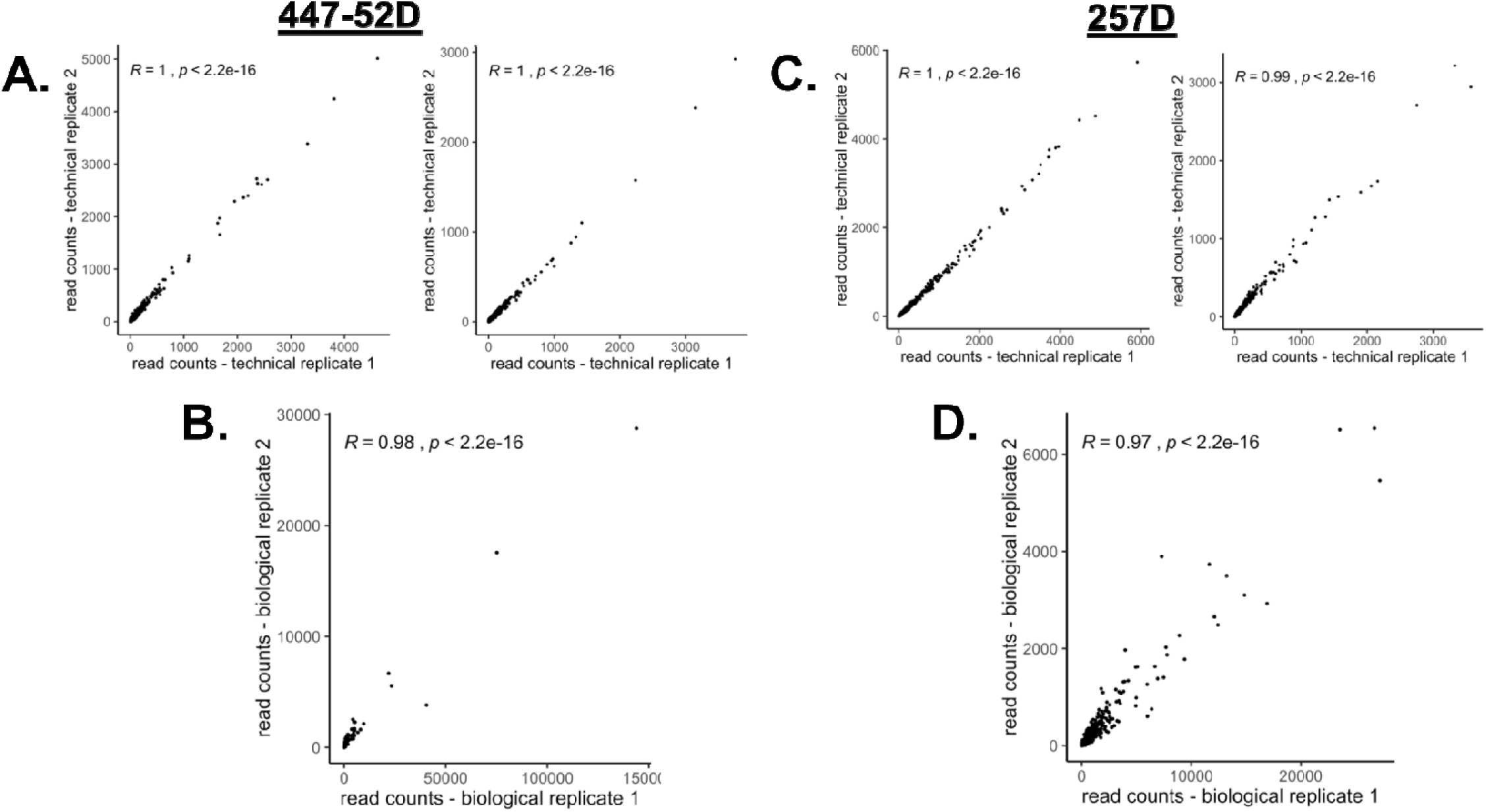
Reproducibility of gp41/V3 Phage-DMS experiments with V3-specific antibodies. Correlation of raw read counts between technical and biological replicates for mAb 447-52D and mAb 257D. **(A, C)** Correlation between replicate wells done in parallel for each experiment. **(B, D)** Correlation between separate experiments performed with gp41/V3 library 1 and 2. Pearson’s correlation is shown at the top.

**Supplemental Figure 4.**
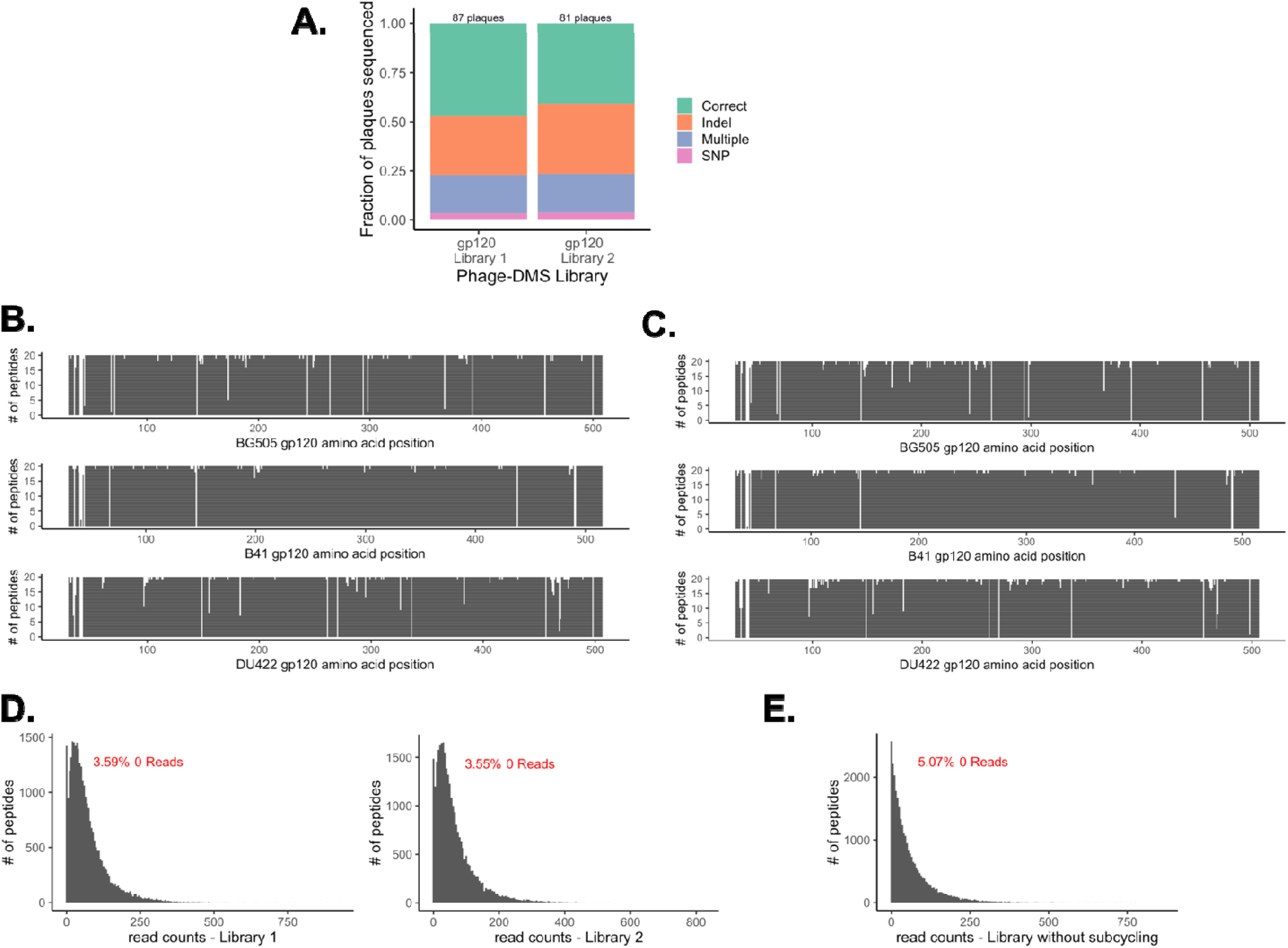
Composition and distribution of sequences in the gp120 Phage-DMS libraries. **(A)** Stacked bar plot depicting results of Sanger sequencing individual plaques from each library. Total number of plaques sequenced is displayed at the top. SNP = single nucleotide polymorphism. **(B)** Coverage of all unique sequences centered at sites across gp120 within gp120 library 1, as determined by deep sequencing at 71-fold coverage. The height of the line at each site corresponds to whether each unique amino acid variant was counted at least once, with 20 total possible. **(C)** Coverage of all unique sequences centered at sites across gp120 within gp120 library 2, as determined by deep sequencing at 65-fold coverage. **(D)** Histogram showing the distribution of all reads sequenced from gp120 library 1 and 2 as determined by deep sequencing. Percent of computationally designed sequences not counted shown in red. **(E)** Histogram showing the distribution of all reads sequenced from a gp120 library amplified without using subcycling PCR. Percent of computationally designed sequences not counted shown in red.

**Supplemental Figure 5.**
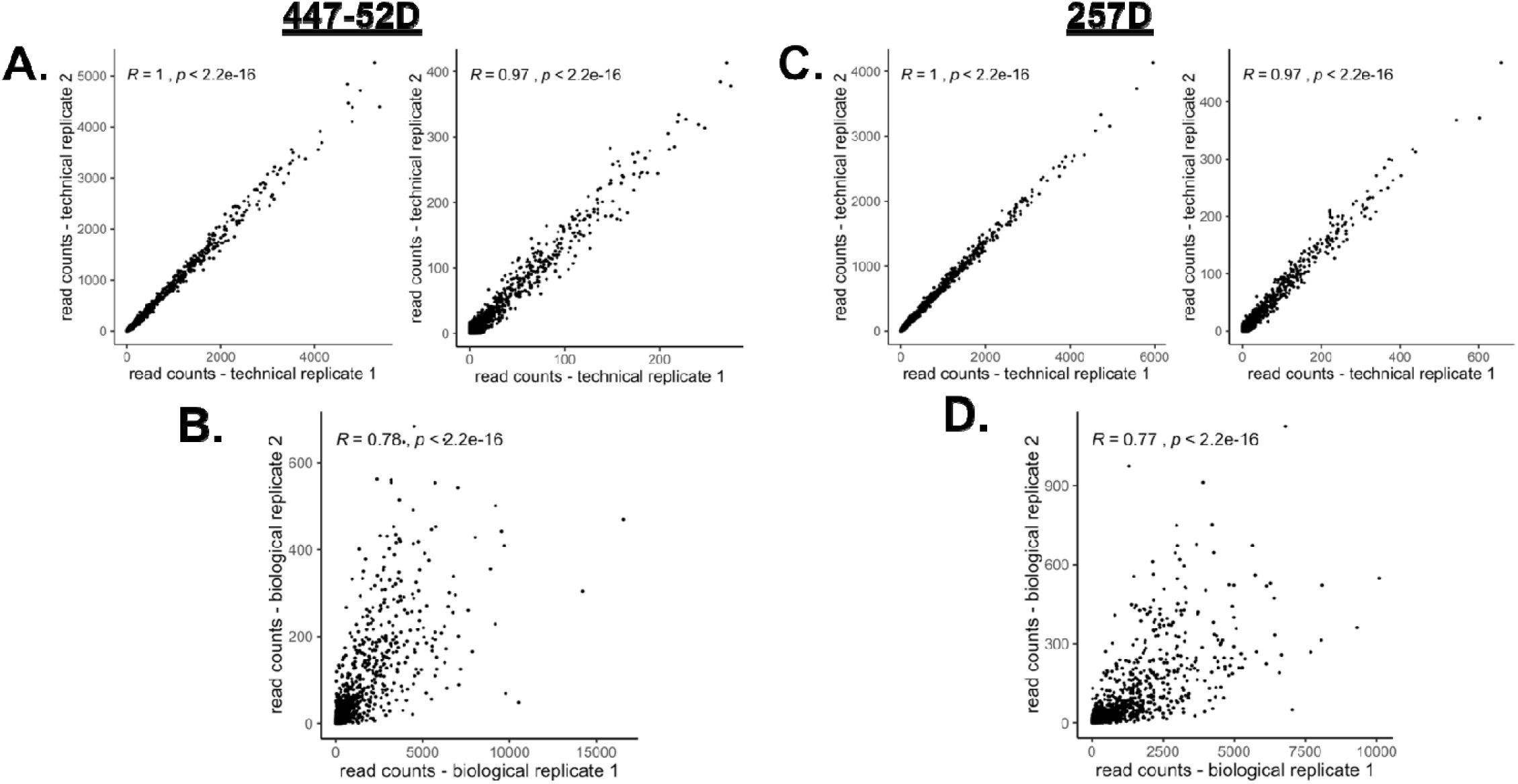
Reproducibility of gp120 Phage-DMS experiments with V3-specific antibodies. Correlation of raw read counts between technical and biological replicates for mAb 447-52D and mAb 257D. **(A, C)** Correlation between replicate wells done in parallel for each experiment. **(B, D)** Correlation between separate experiments performed with gp120 library 1 and 2. Pearson’s correlation is shown at the top.

**Supplemental Figure 6.**
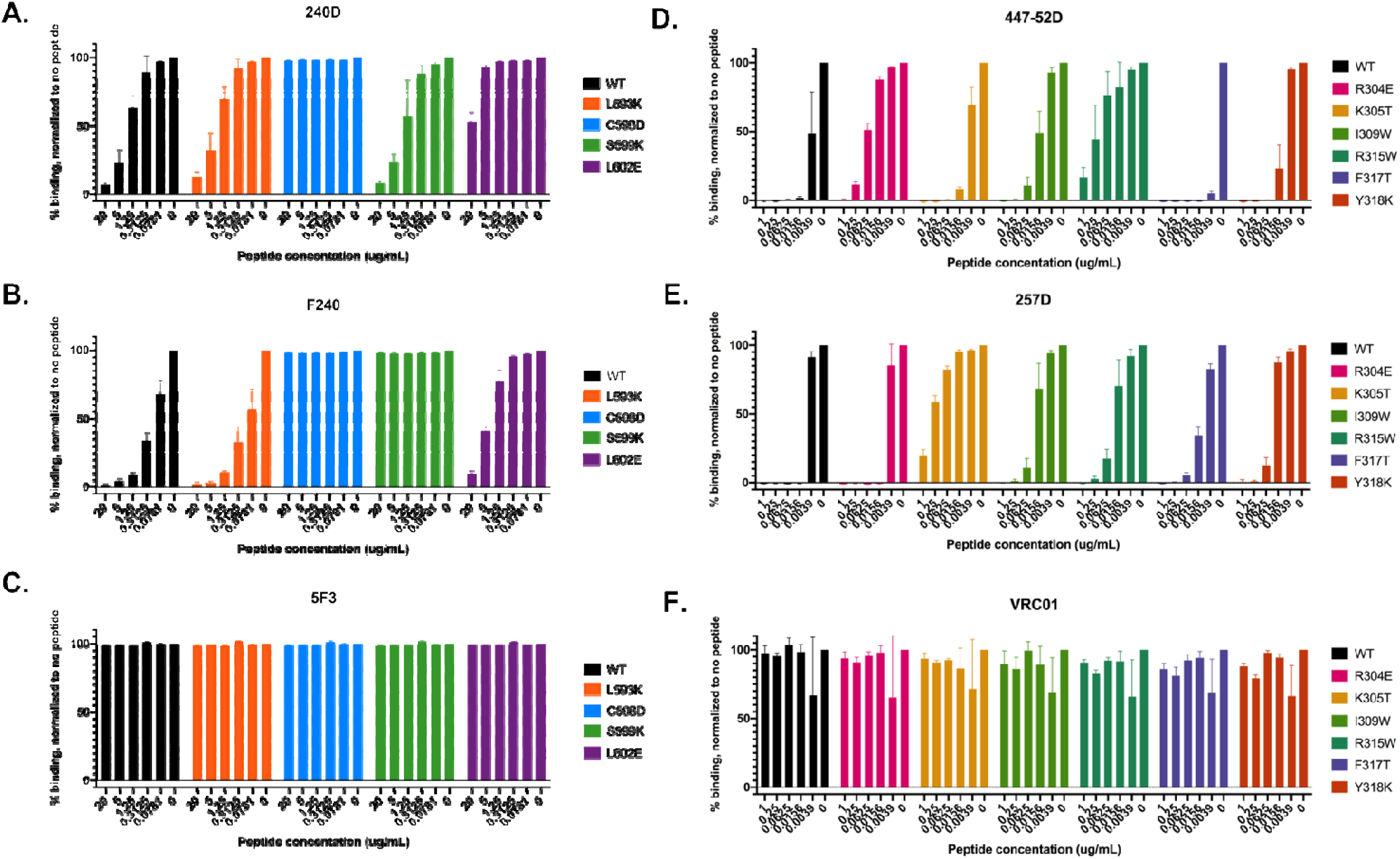
Results of competition peptide ELISAs. **(A-C)** Bar plots show ability of wild type and mutant peptides to block binding of (A) mAb 240D, (B) mAb F240, or (C) mAb 5F3 to MN gp41 protein. 5F3 is a control antibody known to bind to gp41 outside of the C-C loop region. Peptide concentrations are show below each group and represent 4-fold serial dilutions beginning at 20 ug/mL. Gp41-specific antibodies used at a concentration of 0.5 ug/mL, and gp41 protein was coated at 0.5 ug/mL. For reference, the gp41 sequence of MN corresponding to the region the gp41 peptides span is LLGFWGCSGKLICTTTVP. **(D-F)** Bar plots show ability of wild type and mutant peptides to block binding of (D) mAb 447-52D, (E) mAb 257D, or (F) mAb VRC01 to MN gp41 protein. VRC01 is a control antibody known to bind to gp120 outside of the V3 loop region. Peptide concentrations are show below each group and represent 4-fold serial dilutions beginning at 20 ug/mL.V3-specific antibodies used at a concentration of 1 ug/mL, and gp41 protein was coated at 1 ug/mL. For reference, the V3 sequence of SF162 corresponding to the region the V3 peptides span is RKSITIGPGRAFY.

